# Density-based binning of gene clusters to infer function or evolutionary history using GeneGrouper

**DOI:** 10.1101/2021.05.27.446007

**Authors:** Alexander G. McFarland, Nolan W. Kennedy, Carolyn E. Mills, Danielle Tullman-Ercek, Curtis Huttenhower, Erica M. Hartmann

## Abstract

**Motivation:** Identifying gene clusters of interest in phylogenetically proximate and distant taxa can help to infer phenotypes of interest. Conserved gene clusters may differ by only a few genes, which can be biologically meaningful, such as the formation of pseudogenes or insertions interrupting regulation. These qualities may allow for unsupervised clustering of similar gene clusters into bins that provide a population-level understanding of the genetic variation in similar gene clusters.

**Results:** We developed GeneGrouper, a command-line tool that uses a density-based clustering method to group gene clusters into bins. GeneGrouper demonstrated high recall and precision in benchmarks for the detection of the 23-gene *Salmonella enterica* LT2 Pdu gene cluster and four-gene *Pseudomonas aeruginosa* PAO1 Mex gene cluster in 435 genomes containing mixed taxa. In a subsequent application investigating the diversity and impact of gene complete and incomplete LT2 Pdu gene clusters in 1130 *S. enterica* genomes, GeneGrouper identified a novel, frequently occurring *pduN* pseudogene. When replicated *in vivo*, disruption of *pduN* with a frameshift mutation negatively impacted microcompartment formation. We next demonstrated the versatility of GeneGrouper by clustering both distant homologous gene clusters and variable gene clusters found in integrative and conjugative elements.

**Availability:** GeneGrouper software and code are publicly available at https://github.com/agmcfarland/GeneGrouper.

## Background

Physically proximate groups of genes, called gene clusters, are present in many microbial taxa (1). Gene clusters can include genes that form biosynthetic pathways or efflux, secretion, and signaling systems (1–5). Some gene clusters are arranged into one or multiple operons (6). Microbial genomes are under constant gene flux, driven by gene gain, loss, and rearrangements (5,7,8). The identification of intact, conserved gene clusters across different genomes can allow for inferences to be made as to the gene cluster’s functionality, stability, and taxonomic distribution (6,9).

There are different, overlapping approaches used for the identification and classification of gene clusters. Approaches that incorporate a reference database of curated gene clusters include DOOR2, T346Hunter, TADB2.0, and MetaCRAST (10–13). Synteny-based approaches are generally split into those that identify all gene clusters at the genome level compared to within a defined genomic window. Examples of genome-level synteny tools are CSBFinder, GECKO3, and Mauve and genomic window tools include SynFind and SimpleSynteny (14–18). Gene cluster homology search approaches, like MultiGeneBlast or SLING, identify gene clusters that contain all or some of the gene cluster query genes (19,20). After identifying a set of gene clusters, some tools will further aggregate gene clusters into bins using sequence similarity networks or clustering (14,20).

A challenge in analyzing large numbers of gene clusters is that many conserved gene clusters will display little variation in gene content, but that variation may nevertheless be biologically significant, for example an insertion disrupting key genes in a biosynthetic operon (6,21). A population-level understanding of gene cluster content can help to identify which genes are typically located in a gene cluster, and which are variable. Most tools require either custom analysis of tabular outputs or manual inspection of gene cluster synteny plots to identify variations in gene cluster content and their distribution within the analyzed genomes.

We developed GeneGrouper to identify, quantify, contextualize, and visualize the degree of similarity for gene clusters that contain a queried gene of interest in a population of user-supplied genomes. It is designed to work on thousands of genomes and is suitable for use on a personal computer. We demonstrate the utility of GeneGrouper by comparing its unsupervised clustering accuracy with existing tools in the identification of two distinct gene clusters, the 23-gene catabolic microcompartment Pdu gene cluster found in *Salmonella enterica* LT2 and the four-gene MexR/MexAB-OprM Resistance-Nodulation-Division (RND)-type efflux pump gene cluster from *Pseudomonas aeruginosa* PAO1 in 435 taxonomically diverse genomes (22,23). GeneGrouper was next used to examine the diversity and distribution of gene complete and incomplete LT2 Pdu gene clusters in 1130 *S. enterica* genomes. Using GeneGrouper’s visual and tabular outputs, we identify a novel pseudogene present in a subset of otherwise gene-complete LT2 Pdu gene clusters. We replicate the pseudogene *in vivo* and find that it negatively impacts microcompartment formation.

## Implementation

GeneGrouper is written in Python 3 and uses the BioPython and Sci-Kit learn libraries for sequence processing, clustering, and analysis (24,25). Multithreading is implemented via the multiprocessing library (26). GeneGrouper calls on BLAST+, mmseqs2, and MCL for sequence detection, homology searching, and orthology clustering (27–29). Visualizations are generated using R and gggenes (30).

### Input and pre-processing

GeneGrouper requires two inputs: genome files and a translated seed gene sequence. Genome files must be in GenBank file format like those from the NCBI Refseq database (31). All genome files have coding sequence features extracted and stored in an SQLite database. A BLAST database is constructed from all extracted amino acid sequences.

### Seed homology searching

A BLASTp search for the translated seed gene is performed using user-specified identity and coverage thresholds **(Fig. 1A i)**. Upstream and downstream genomic regions of lengths corresponding to user-specified base-pair distances are extracted. In instances where extracted regions overlap, the region with the highest E-value is chosen. The genomic position of hits and the amino acid sequences within the defined genomic region are written to a seed gene-specific SQLite database. All sequences within the defined genomic region are extracted and stored as a FASTA file.

**Figure 1:**
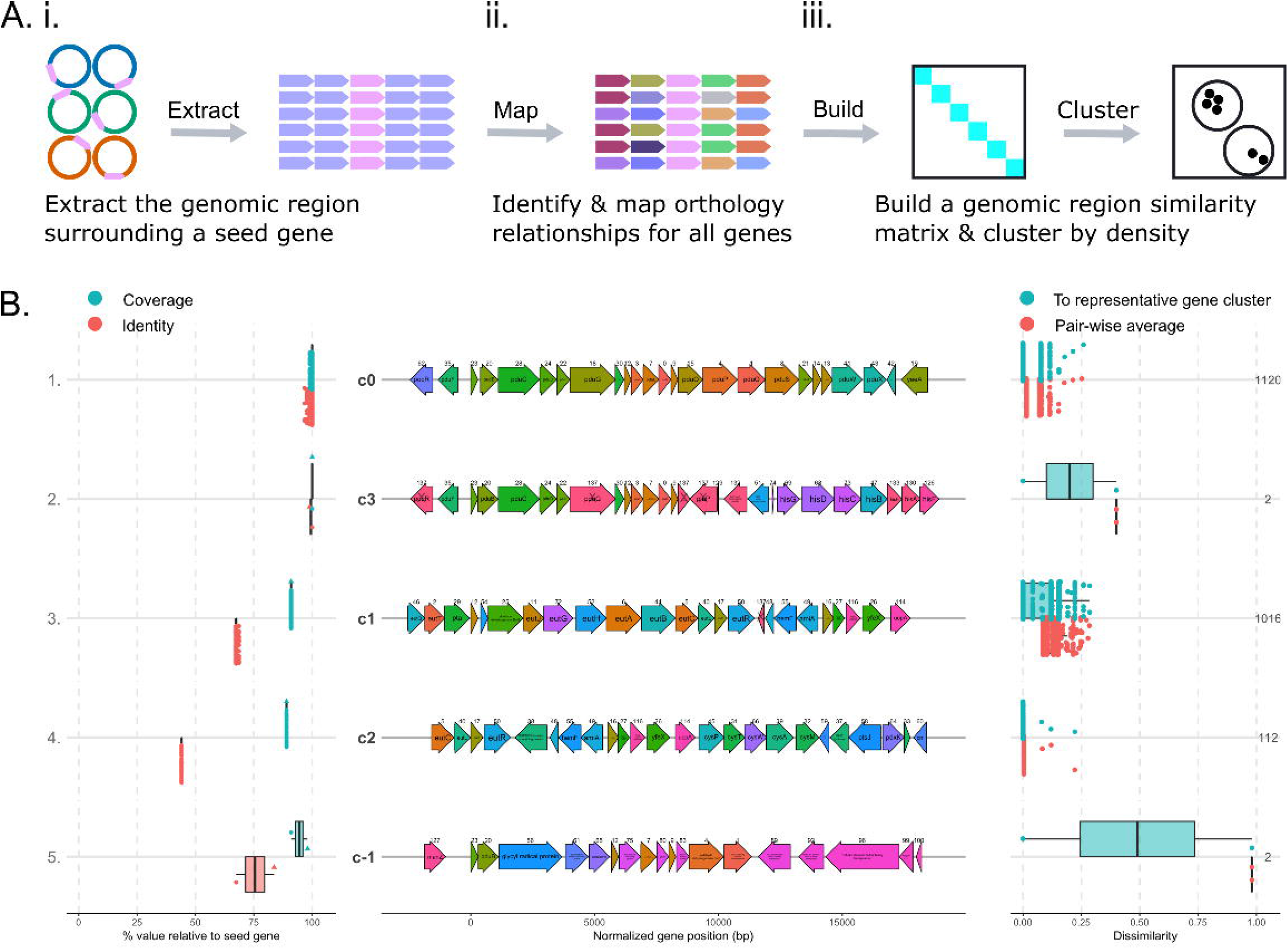
**A (i-iii).** Summarized gene extraction and clustering workflow. **B.** GeneGrouper’s main output for a search of Pdu gene clusters in 1130 S. enterica genomes. The left panel displays BLAST hit statistics for each seed gene belonging to a gene cluster in a cluster label. The middle panel displays the gene cluster architecture that is representative of the cluster label. The right panel shows the dissimilarity of other gene clusters in the cluster label. ‘X’ indicates a pseudogene. Numbers above gene arrows are internal orthology identifiers.

### Orthology inference and assignment

GeneGrouper assigns orthology to all sequences extracted from the defined genomic region using an internal pipeline. The internal orthology identification scheme takes as input a FASTA file generated during the pre-processing phase containing all detected amino acid sequences. Sequences are clustered using mmseqs2 linclust to generate a set of proximate orthology relationships, producing a set of representative amino acid sequences in FASTA format. An all-vs-all BLAST search is performed, with the resulting hits table filtered for identity, coverage, E-value, and desired number of matches. The E-values from the filtered hits table is used as an input for Markov Graph Clustering with MCL. MCL is run over multiple inflation values, with the lowest inflation value containing the highest count of unique orthologs selected by default. The MCL and mmseqs2 linclust ortholog group assignments are transferred to every sequence and stored **(Fig. 1A ii)**.

### Genomic region clustering

Pairwise Jaccard distances are calculated for all genomic regions **(Fig. 1A iii)** (25). The DBSCAN algorithm is then run on the resulting dissimilarity matrix using a fixed minimum cluster size value over increasing epsilon values (32). For each epsilon value, the number of clusters, noise, silhouette score, and Calinksi-Harabasz score are calculated (33). The epsilon value demonstrating the best separation of clusters (defaulting to the highest Calinksi-Harabasz score) is selected. The previously constructed Jaccard distance matrix is subsetted for regions within each DBSCAN cluster label. For each cluster label, the mean dissimilarity for each region is calculated. The region with the lowest mean dissimilarity is selected as the representative for that cluster label.

### Outputs

Tabular outputs containing the cluster label, region identifier, mean cluster label dissimilarity, and relative dissimilarity to the cluster representative is generated. Each unique cluster is assigned a numeric label. All gene regions that could not be assigned to any cluster are grouped into cluster label ‘c-1’. Three main visualizations are produced: The representative gene regions for each cluster with gene annotations **(Fig. 1B)**, the dissimilarity contained within each gene region, and the unique variants found in each gene region along with their count. Users can query specific clusters and generate a fourth visualization type showing the count, dissimilarity, and structure of each unique gene cluster within that queried cluster.

## Results and Discussion

### Gene cluster selection and rationale

435 genomes with chromosome-level assemblies were downloaded from the NCBI Refseq database on March 23 2021 (31). These genomes belonged to six taxa: *Salmonella enterica, Klebsiella pneumoniae, Pseudomonas aeruginosa, Citrobacter, Enterobacter*, and *Clostridium* (**Table S1**). These genomes were searched for the presence of complete and partial LT2 Pdu and PAO1 Mex gene clusters (**Table 1)**. The LT2 Pdu gene cluster contains 23 genes encoding bacterial microcompartments (BMC) that allow for the metabolism of 1,2-propanediol. The LT2 Pdu gene cluster was selected to test whether GeneGrouper could detect and accurately bin a large gene cluster that contains multiple paralogs (i.e., *pduA, pduJ*, and *pduT*), present in multiple phylogenetically distinct genomes (i.e., *S. enterica, K. pneumoniae*, and *Citrobacter*) while avoiding the inclusion of other separate microcompartment gene clusters present in all six genera that share some orthologs (9). The PAO1 Mex gene cluster (*mexR, mexA. mexB*, and *oprM*) encodes for an RND efflux pump that has multiple homologs present within a genome and across virtually all Gram-negative species (2). The PAO1 Mex gene cluster is distinguished by its MarR-type proximal repressor, MexR, in *P. aeruginosa*. The PAO1 Mex gene cluster was selected to test whether GeneGrouper could specifically detect a short gene cluster with multiple homologs within a species, and across all five Gram-negative taxa in our collection of genomes.

**Table 1:**
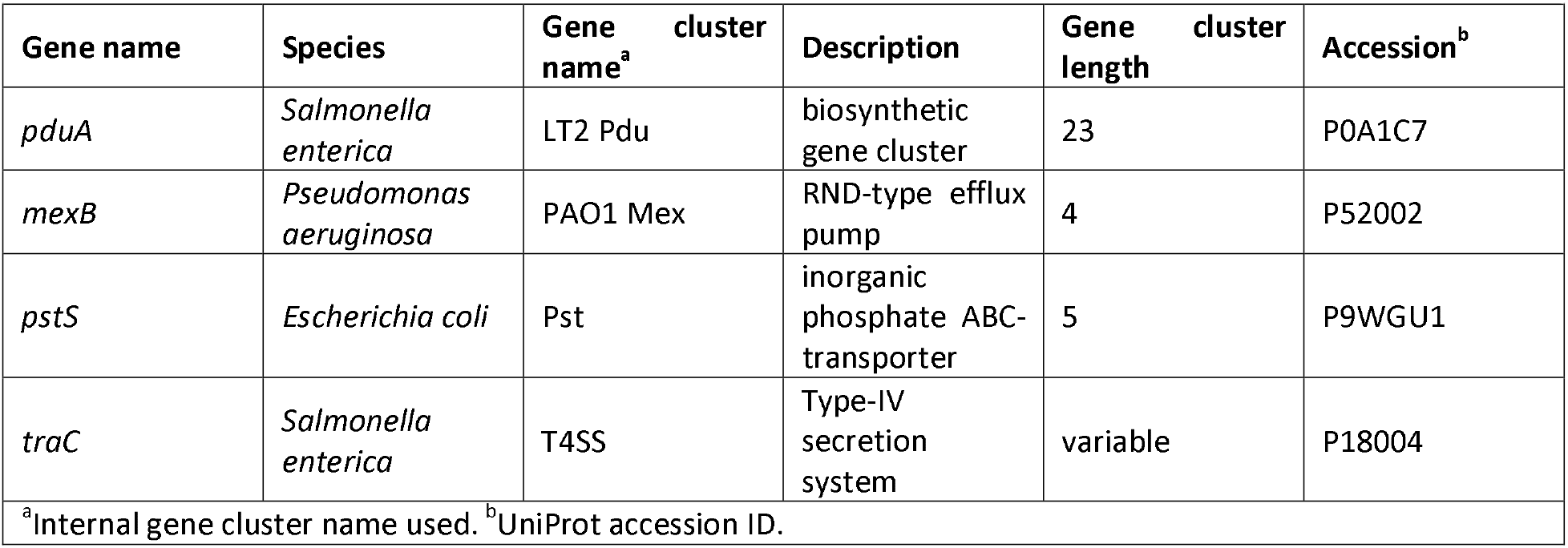
Genes used for searches and comparisons, and the gene clusters they represent.

### Identification of full or partial Pdu and MexAB-OprM gene clusters using different tools

We compared the capacities of GeneGrouper, MultiGeneBlast, and CSBFinder to detect full or partial LT2 Pdu and PAO1 Mex gene clusters in all 435 genomes. Each tool was limited to only detecting gene clusters, and no internal scoring or clustering algorithm was used. Each individual gene cluster was then scored in a standard manner as ‘full’ (100% of expected gene clusters present), ‘partial’ (<100-70% present), or ‘other’ (<70% present). In this manner, the capacity for different tools to detect gene clusters was standardized to allow for direct comparison of the detection of specific gene clusters and limit idiosyncrasies in clustering/scoring approaches.

For the detection of the LT2 Pdu gene cluster, GeneGrouper was run using the translated *S. enterica* LT2 PduA sequence as a seed, and a genomic search space of 2,000 bp downstream and 18,000 bp upstream to capture the two genes downstream and 22 genes upstream of *pduA*. PduA was selected as the seed because it is a member of the pfam00936 protein family, which is the hallmark indication of BMC loci (9). For the detection of the PAO1 Mex gene cluster, the *P. aeruginosa* PAO1 MexB sequence was used as a seed with a uniform search space of 10,000 bp upstream and downstream. For both searches a ≥60% identity and ≥80% coverage threshold was used. The orthology assignments for each gene in the Pdu or PAO1 Mex gene clusters were then used to score gene clusters.

MultiGeneBlast was run in search mode with default settings on each individual genome using an input FASTA file that contained all the translated gene sequences belonging to either LT2 Pdu or PAO1 Mex gene clusters. BLAST results for each identified gene cluster were filtered such that each individual query gene was matched to its single best hit, with a coverage cutoff of ≥80%, and no identity cutoff to allow for phylogenetically distant hits to be preserved. Each gene cluster was then scored.

CSBFinder inputs were pre-processed prior to gene cluster searching. The proteomes for all six genera were generated by clustering with mmseqs2 linclust (34). Afterwards, orthology identification was performed using OrthoFinder with default settings (35). Genomes with orthology assignments were then converted into the CSBFinder format. The orthology assignments for each gene present in either the LT2 Pdu or PAO1 Mex gene clusters were converted to a ‘patterns’ file and used to search all genomes for the respective gene clusters and then scored.

All approaches identified full LT2 Pdu gene clusters in almost all S. *enterica* and *Citrobacter* sp, and most *K. pneumonia* genomes. Between all three approaches, 224 to 288 full and 13 to 20 partial LT2 Pdu gene clusters were predicted **(Fig. 2A**). CSBFinder had the most conservative results and did not identify any *K. pneumoniae* genomes carrying the LT2 Pdu gene cluster. GeneGrouper and MultiGeneBlast had comparable counts for full and partial LT2 Pdu gene cluster detection, with GeneGrouper identifying fewer full and more partial LT2 Pdu gene clusters.

**Figure 2:**
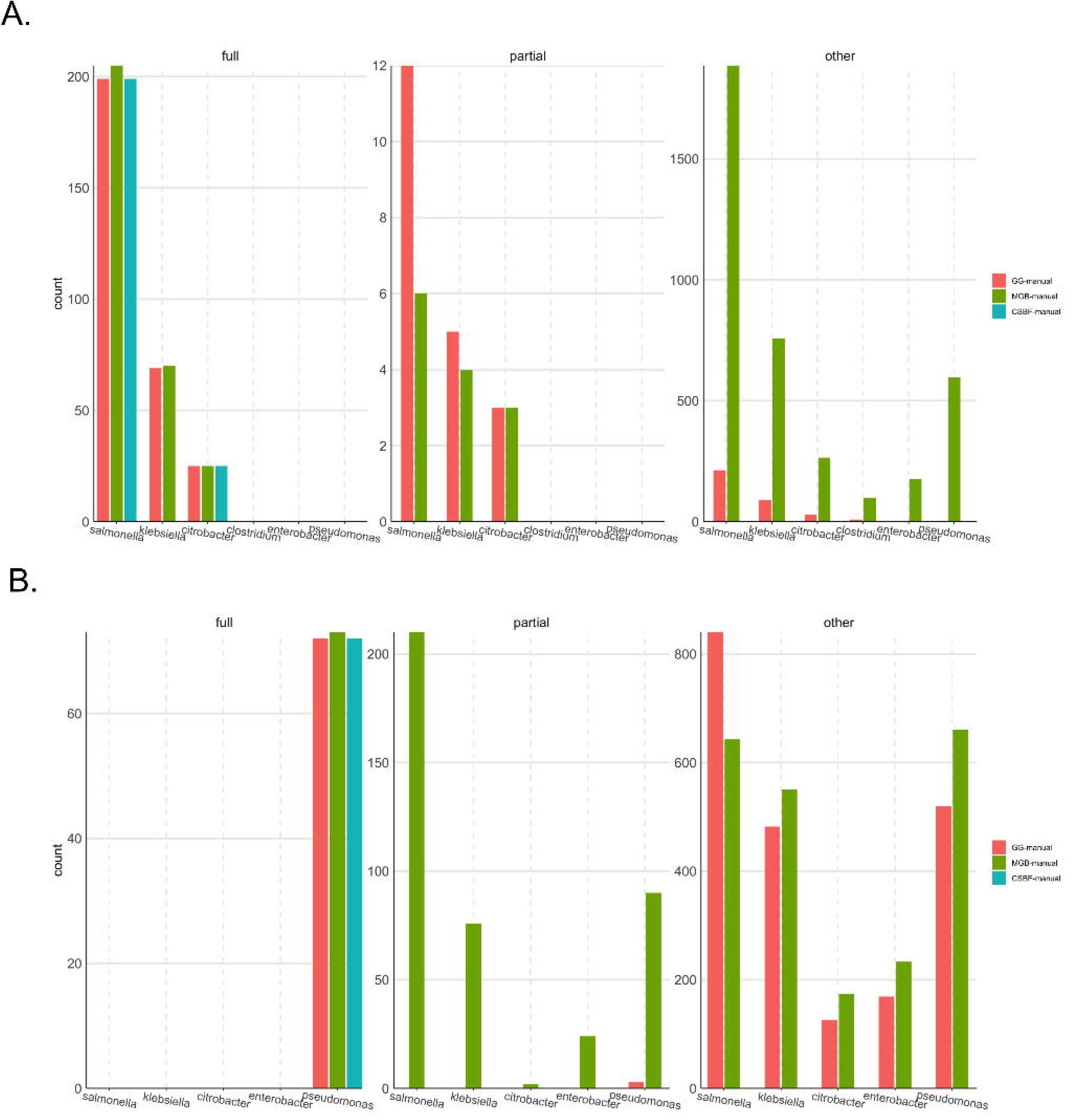
Counts of gene cluster types. **A.** LT2 Pdu. **B.** PAO1 Mex.

We next searched the same set of genomes for the presence of the PAO1 Mex efflux pump operon and its proximal regulator, MexR. All approaches identified either 72 or 73 genomes carrying the full PAO1 Mex gene cluster, and all were in *P. aeruginosa* (**Fig. 2B)**. CSBFinder only identified full MexAB-OprM gene clusters. GeneGrouper iden2tified three partial PAO1 Mex gene clusters, all in *P. aeruginosa*. MultiGeneBlast identified 402 partial PAO1 Mex gene clusters, distributed throughout *P. aeruginosa*, *S. enterica, K. pneumoniae*, *Citrobacter*, and *Enterobacter*.

Using our standardized methods for the detection of either the LT2 Pdu or PAO1 Mex gene cluster, all tools compared similarly in the capacity to detect genomes carrying a full gene cluster. CSBFinder reported lower numbers of full or partial gene clusters in phylogenetically distant genomes. This is likely due to the orthology pre-processing step using OrthoFinder. MultiGeneBlast had higher numbers of partial gene clusters detected, especially for PAO1 Mex gene clusters. The large number of partial gene clusters was likely due to the BLAST-based scoring system that did not use an identity cutoff, compared to the reference orthology group approach used by GeneGrouper and CSBFinder. Importantly, these results demonstrate that GeneGrouper detects similar numbers of full or partial gene clusters compared to existing methods using a standardized scoring method.

### Accuracy of GeneGrouper automated gene cluster binning

GeneGrouper uses an unsupervised learning approach to aggregate each individual gene cluster into a cluster label. Each cluster label should contain gene clusters that have similar, but not always identical, gene content, over a defined distance. Therefore, a cluster label will likely contain both full and partial gene clusters but should not contain unrelated gene clusters. We tested whether this was the case by comparing the results of our standardized gene cluster identification with the clustering results produced by GeneGrouper. Prior to testing, the ground truth of each genome for the presence of a full, partial, or absent LT2 Pdu/ PAO1 Mex gene cluster was determined.

The LT2 Pdu gene cluster was searched for in all genomes with GeneGrouper using the same parameters as before. GeneGrouper assigned 654 different gene clusters to four different cluster labels and had 12 gene clusters with no clustering solution that were placed in cluster label ‘c-1’ **(Table 2, Fig. S1, Fig. S2A)**. Cluster label ‘c0’ contained the Pdu gene cluster from *S. enterica* LT2 and was designated as being the cluster label that contained all expected instances of full or partial LT2 Pdu gene clusters (from here on referred to as GG-cluster). Overall, the precision and recall scores of GG-cluster compared favorably with the scores from the standardized approaches **(Fig. 3A)**. GG-cluster had a lower precision when used to predict the presence of only full LT2 Pdu gene clusters. However, the precision increased to 1 when predicting full or partial Pdu gene clusters. Comparatively, the recall remained almost unchanged, with a score of 1 when identifying full Pdu gene clusters and 0.99 when identifying full or partial Pdu gene clusters. GG-cluster missed one instance of a partial Pdu gene cluster that was assigned to cluster label ‘c-1’, which contains gene clusters for which no clustering solution was found. This Pdu cluster was split in two in the referenced assembly, being present at the start and end of different contigs. These results demonstrate that GG-cluster accurately identified almost all LT2 Pdu gene clusters of either full or partial status and did not incorrectly identify non-Pdu gene clusters as LT2 Pdu gene clusters.

**Figure 3:**
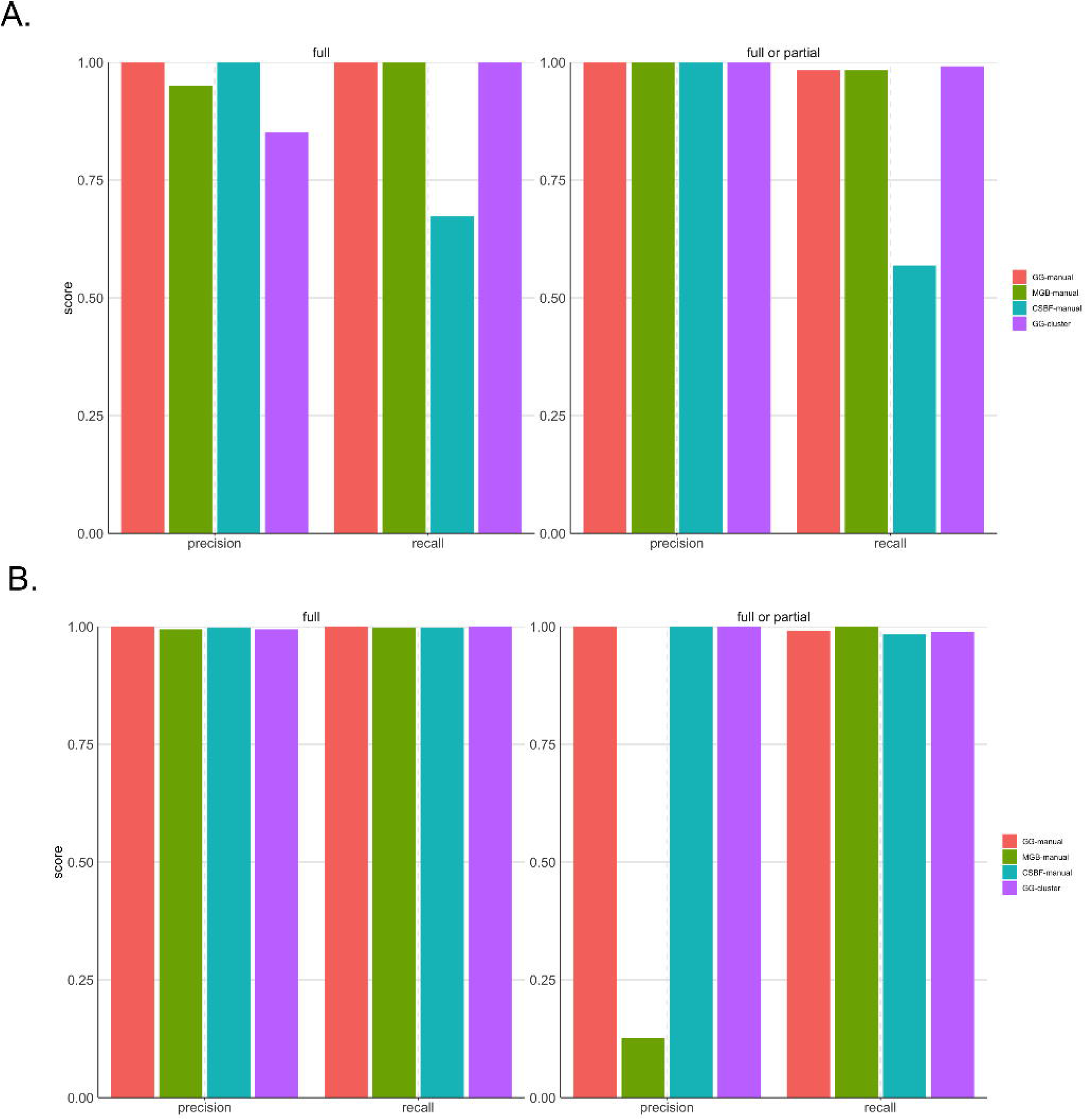
Precision and recall scores for comparisons. **A.** LT2 Pdu. **B.** PAO1 Mex.

**Table 2:** Search parameters used for GeneGrouper in this study.

| **Table 2** |  |  |  |  |  |  |  |  |
| --- | --- | --- | --- | --- | --- | --- | --- | --- |
| **Gene cluster name<sup>a</sup>** | **Upstream-Downstream search length (bp)** | **Seed identity (%) - Coverage (%) - Hitcount** | **Total gene clusters found** | **Total cluster labels** | **Total unclustered gene clusters** | **Genomes with hit** | **Dataset** | **Run time (h:m:s)<sup>b</sup>** |
| LT2 Pdu | 2000-18000 | 30 - 80 - unlimited | 654 | 4 | 12 | 324 | 435 mixed | 00:01:48 |
| PAO1 Mex | 10000-10000 | 30 - 80 - unlimited | 2213 | 40 | 55 | 423 | 435 mixed | 00:03:22 |
| Pst | 8000-8000 | 15 - 70 - 1 | 394 | 7 | 1 | 394 | 435 mixed | 00:00:54 |
| T4SS | 20000-20000 | 15 - 70 - unlimited | 81 | 4 | 10 | 59 | 435 mixed | 00:00:37 |
| LT2 Pdu | 2000-18000 | 30 - 80 - unlimited | 2252 | 4 | 2 | 1128 | 1130 single | 00:04:12 |
| <sup>a</sup>Internal gene cluster name used. <sup>b</sup>hours:minutes:seconds |  |  |  |  |  |  |  |  |

PAO1 Mex gene clusters were searched for using GeneGrouper as previously described, identifying 2213 gene clusters contained within 40 cluster labels (**Table 2, Fig. S3, Fig. S2B**). Cluster label ‘c0’ contained the *P. aeruginosa* PAO1 Mex gene cluster and was designated as being the cluster label that contained all expected instances of full or partial PAO1 Mex gene clusters and thus designated as GG-cluster. All approaches had between 0.99 and 1 precision and recall for the identification of full PAO1 Mex gene clusters (**Fig. 4B)**. GG-cluster missed assigning one PAO1 Mex gene cluster that was binned in cluster label ‘c-1’ and missed three instances where MexB was a pseudo gene and therefore could not be detected in the initial search. For the prediction of full or partial PAO1 Mex gene clusters, MultiGeneBlast had a precision score of 0.12, likely due to the high degree of sequence identity between homologous RND efflux pump components. All other tools scored between 0.98 and 1 for precision and recall. These results indicate the GeneGrouper clustering assignment can sort through different variations of highly similar gene content and identify specific RND efflux pump components.

**Figure 4:**
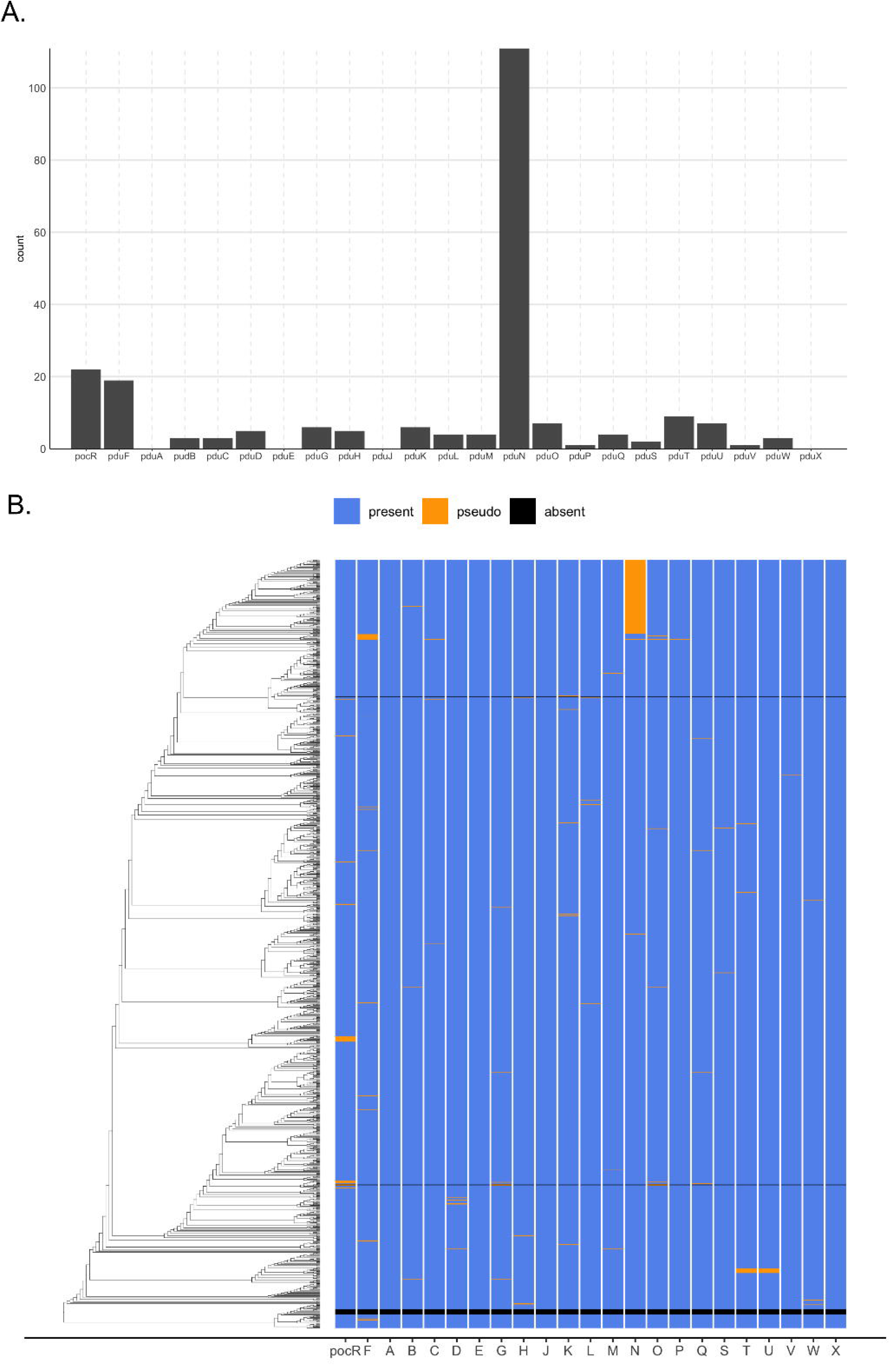
Whole genome analysis of 1130 *S. enterica* genomes for the LT2 Pdu gene cluster. **A.** Count of pseudo gene occurrence. **B.** Whole-genome phylogeny (left) and a presence/absence/or pseudo gene matrix for the LT2 Pdu gene cluster.

### Application: Distribution and diversity of full and partial Pdu gene clusters in *S. enterica*

Although *S. enterica* is known to carry the LT2 Pdu gene cluster, it is unclear how full and partial LT2 Pdu gene clusters are distributed within the species, and whether unique insertions, deletions, or outright losses of the gene cluster have occurred and propagated. This is of interest because even in the presence of interruptions, functional LT2 Pdu gene cluster variants may still exist, and can inform on which genes may not be necessary. We used GeneGrouper to search for the LT2 Pdu gene cluster in in 1130 complete or chromosomal-level genome assemblies from the RefSeq database downloaded on March 23, 2021 (**Table S2**) (30). The *S. enterica* LT2 PduA sequence was used as a seed to search and cluster the gene content for a genomic region of 2000 bp downstream and 18000 bp upstream of any *pduA* homolog (**Table 2**). The search returned four distinct cluster labels with distinct gene clusters and two total unclustered gene clusters, which were visualized using GeneGrouper’s visualization command **(Fig. 1B).** GeneGrouper reports the Jaccard dissimilarities of each region within a cluster relative to the region representative so that differences in gene content can be efficiently quantified and assessed. Cluster label ‘c0’ contained the *S. enterica* LT2 strain LT2 Pdu gene cluster, which had zero dissimilarity with the representative region of cluster label ‘c0’. In total, cluster label ‘c0’ contained 1120 regions with a 0 and 0.076 dissimilarity at the 50th and 95th percentiles, respectively. These low dissimilarities indicated that cluster label ‘c0’ had very little variation in gene content relative to its representative region.

To examine the variability in gene content within cluster label ‘c0’, GeneGrouper’s cluster inspection command was run to visualize the count of identical occurrences of each gene cluster **(Fig. S4).** 48 separate identical gene clusters were present, the majority of which had all 23 LT2 Pdu genes. The tabular output was queried to reveal that of the gene clusters identified, 920 (81.4%) carried all 23 LT2 Pdu genes, 10 (0.88%) did not have a LT2 Pdu gene cluster identified, and the remaining 200 (17.6%) had predicted LT2 Pdu gene clusters with between one and five pseudogenes. Interestingly, gene clusters carrying a *pduN* pseudogene but otherwise complete were the most common non-complete gene cluster observed (**Fig. 4A)**. A whole-genome phylogeny of all 1130 genomes was created using Phylophlan 3.0 with the provided 400 marker sequence database and visualized with ggtree along with a presenceabsence matrix of each Pdu component extracted from GeneGrouper’s tabular output (36,37). We found that genomes with *pduN* pseudogenes were present almost entirely in the same section of the phylogenetic tree (**Fig. 4B).** This is a surprising finding, as PduN is a necessary component for proper Pdu microcompartment formation (22). PduN is a member of the BMC vertex protein family (pfam03319), which are necessary for capping the vertices of BMCs and imparting the standard polyhedral morphology (22,38,39). Absence of PduN leads to malformed and elongated microcompartment structures and disrupted growth on the substrate 1,2-propanediol (22). The PduN mutation found in strain *S. enterica* Ty2 (GCF_000007545.1) contained a nucleotide deletion at position 68 that resulted in a frame-shift mutation **(Fig. S5)**.

The effects of this deletion on microcompartment formation were tested in *S. enterica* LT2 **(See Supplemental Text for methods)**. In order to determine the effect of the PduN frameshift and resulting pseudogene seen in our analysis, strains containing this frameshift (denoted ΔN::N*) were generated and compared to strains containing the intact Pdu gene cluster (WT), a full PduN deletion (ΔN), and a negative control lacking the essential pfam00936 genes *pduA* and *pduJ* (ΔAΔJ) **(Fig. 6)** (40). Microcompartment formation was tested using a GFP encapsulation assay, in which GFP is targeted to microcompartments using an N-terminal signal sequence sufficient for microcompartment targeting (41,42). We found that strains expressing the *pduN* pseudogene (ΔN::N*) exhibited aberrant microcompartment morphologies similar to those observed in the *pduN* knockout strain (ΔN), indicating improper microcompartment assembly due to a loss of vertex capping. This phenotype is distinct from the bright fluorescent puncta throughout the cytoplasm in the WT strain, indicative of normal microcompartment assembly and morphology, as well as the ΔAΔJ negative control containing polar bodies, indicative of aggregation. These results demonstrate the utility of GeneGrouper in rapidly identifying pseudogenes that dramatically alter the functionality of BMC gene clusters.

### Additional gene cluster searches using GeneGrouper

To demonstrate the applicability of GeneGrouper to other gene cluster types and use cases, we searched for an additional two different seed genes (**Table 1**). The Pst gene cluster (*pstSCAB*) is present in many Gram-negative and positive bacteria, encodes for a four-component phosphate ABC-transporter and is adjacent to the negative phosphate regulon regulator, *phoU* (43). The Pst gene cluster is present in many Gram-negative and positive bacteria and regulates the uptake of inorganic orthophosphate (44). We wanted to test whether distant or proximal homologs of the *Escherichia coli* Pst gene cluster were present in our genomes. Identity and coverage parameters were lowered to 15% and 70%, respectively, and only the best BLAST hit from each genome was kept (**Table 2**). A total of 393 gene clusters were binned into six cluster labels **(Fig. S6A,B)**. Interestingly, *S. enterica, K. pneumoniae*, and *Enterobacter* had similar gene cluster arrangements, even between clusters, with the main difference being the genes upstream of the Pst gene cluster. Interestingly, the Pst gene cluster has been described in *Clostridium* and verified to be a homolog of the *E. coli* Pst gene cluster (43). In this search, only one *Clostridium* genome has a Pst gene cluster identified and placed alone in cluster label ‘c-1’, suggesting that the Pst gene cluster may not be carried by all *Clostridium*. Another unexpected finding was that only 62.8% of *P. aeruginosa* genomes carried a homolog of *pstS* in a conserved gene cluster assigned to cluster label ‘c0’. However, gene clusters in this cluster label lacked other members of Pst and instead were associated with type II secretion genes. This context suggests that the *pstS* gene in cluster label ‘c0’ may serve a different functional role compared *pstS* found in the Pst gene cluster in *E. coli* (45,46).

In another example use case, *traC*, a type IV secretion system (T4SS) gene found in integrative and conjugative elements (ICEs) was searched for in all 435 genomes (**Table 2)** (47). ICEs have highly variable gene content across both its cargo genes and the components necessary for integration and conjugation (48). To search for *traC*, identity and coverage parameters were maintained as above, with unlimited numbers of hits per genome. A 20000 upstream/downstream genomic range was used, which is on the lower end of ICE sizes (shown to range in size from 37-143 kb) (49). Clustering returned 74 separate gene clusters binned in four different cluster labels, and 10 gene clusters assigned to cluster label ‘c-1’ **(Fig. S7A,B)**. Expectedly, there was high dissimilarity within cluster labels. However, one cluster label, ‘c3’, exhibited low mean dissimilarity and was present in both *S. enterica* and *K. pneumoniae* genomes, suggesting these genomes carry the same ICE. Cluster label ‘c0’ was found in 46% of all *K. pneumoniae* genomes, raising the possibility of a particular mobile or ancient ICE acquisition.

## Conclusions

We demonstrate that GeneGrouper is a simple and accurate tool for identifying gene clusters of interest in a large number of genomes using a single seed gene and a specified genomic window. The use of a gene cluster representative for each cluster label allows for a more intuitive understanding of the diversity of gene clusters that exist in a population, especially when coupled with additional visual metadata, and allow for easy identification and comparison of biologically relevant features. The provided tabular outputs allows for researchers to further probe identified gene clusters for their own specific questions. There exist some limitations in our approach, namely the absence of gene clusters that do not have the seed gene and the presence of incomplete gene clusters due to low-quality assembly genomes. Despite these limitations, comparisons with existing gene cluster detection tools demonstrates that GeneGrouper’s automated clustering and overall approach provides similarly accurate predictions.

In an example application, GeneGrouper was used to determine whether the LT2 Pdu gene cluster was present in 1130 complete *S. enterica* genomes and, if so, how intact the gene cluster was. We further probed the consequences of pseudogene formation in *pduN* and found that the *pduN* pseudogene results in formation of distinct, aberrant microcompartment structures similar to those observed in a *pduN* knockout strain. The example application demonstrates that GeneGrouper enables researchers to rapidly identify gene clusters containing unusual features, specifically pseudogenes that may disrupt proper function, by contextualizing their occurrence against other highly similar gene clusters in a population. This comparative approach has the potential to save time in situations where researchers are choosing model gene clusters for a study by identifying common and unusual related gene clusters. This can help to prevent erroneous conclusions if studies are performed on a gene cluster containing unique genetic features compared to the typical gene cluster in a population.

**Figure 5:**
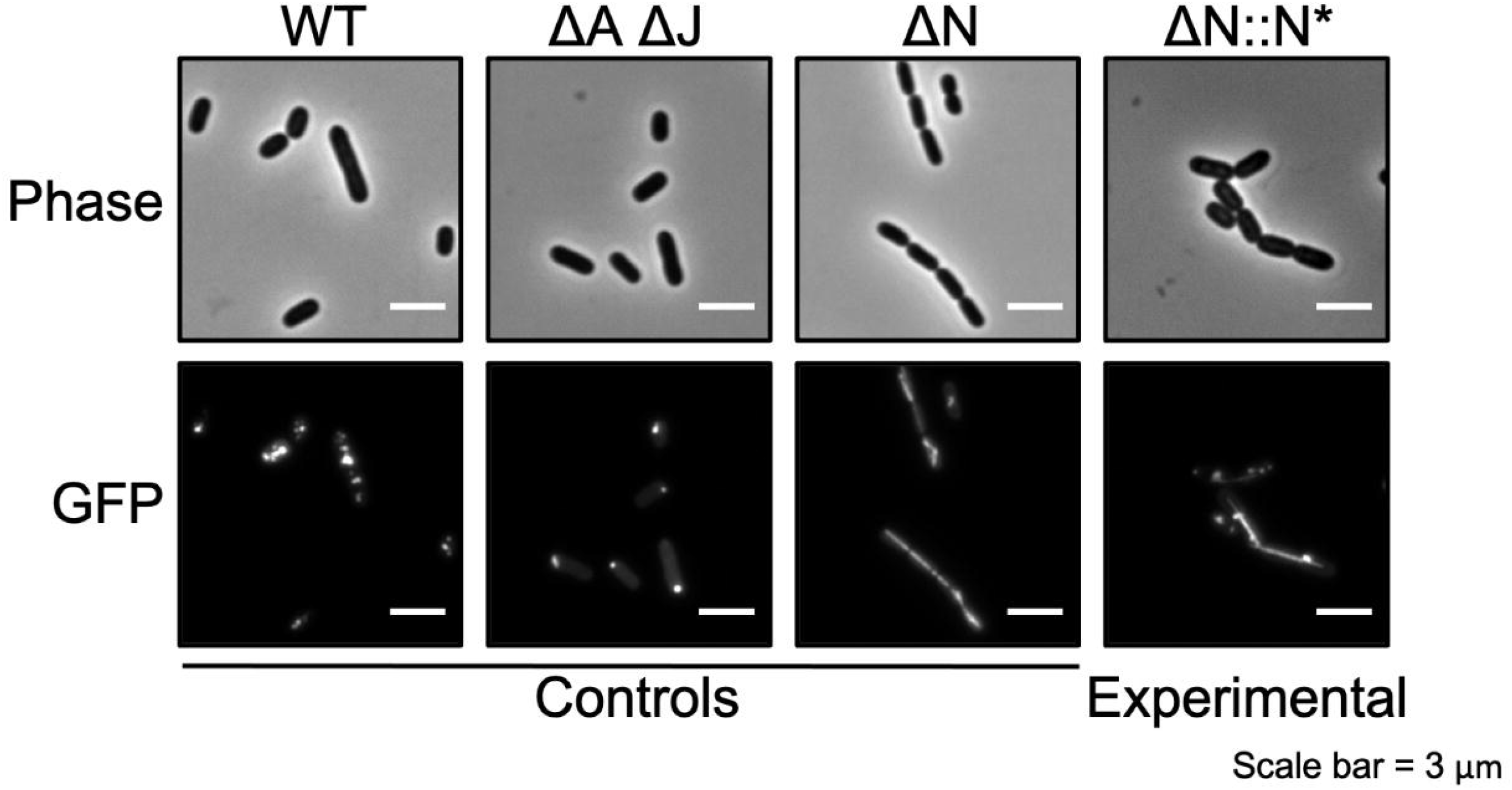
Phase contrast and GFP fluorescence microscopy images of various Salmonella strains expressing ssD-GFP (GFP tagged with the signal sequence from PduD). Row labels indicate micrograph type and column labels indicate bacterial strain. GFP fluorescence images depict fluorescent, cytosolic puncta indicative of microcompartments in the wild type (WT) strain. Fluorescent polar bodies in the *pduA pduJ* double knockout strain (ΔA ΔJ) indicate improper compartment formation. Elongated fluorescent structures in the pduN knockout (ΔN) and *pduN* frameshift (ΔN::N*) indicate improper vertex capping.

## Acknowledgments

Experimental research was supported by the Searle Leadership Fund (E. M. H.), the Biotechnology Training Program (A. G. M.), the Army Research Office (grant W911NF-19-1-0298 to D. T. E.), and the National Science Foundation Graduate Research Fellowship Program (grant DGE-1842165 to N. W. K).

---

**Figure S1:** GeneGrouper main output for the LT2 Pdu gene cluster after searching 435 genomes.

**Figure S2:** GeneGrouper heatmap output displaying the percentage of genomes searched that have at least one gene cluster in a cluster label. Asterisks indicate that a genome had more than one gene cluster in a single cluster label. **A.** LT2 Pdu gene cluster. **B.** PAO1 Mex gene cluster.

**Figure S3:** GeneGrouper main output for the PAO1 Mex gene cluster after searching 435 genomes.

**Figure S4:** GeneGrouper cluster inspection output of cluster label ‘c0’ from Figure 1. Counts of each unique gene cluster architecture (termed subcluster) (left panel), gene cluster architecture representative (middle panel), and dissimilarity to the first subcluster (right panel). ‘X’ indicates a pseudogene. Numbers above gene arrows are internal orthology identifiers.

**Figure S5:** Alignment of *pduN* and the pduN pseudogene. Pair-wise sequence alignments of the translated and untranslated *pduN* frameshifted pseudogene (N_pseudo) and the wild type *pduN* translated and untranslated sequence (N_real)

**Figure S6:** GeneGrouper output for the Pst gene cluster after 435 genomes. **A.** Main output. **B.** Heatmap output.

**Figure S7:** GeneGrouper output for the T4SS gene cluster after 435 genomes. **A.** Main output. **B.** Heatmap output.

## Supplemental methods

Methods used to generate knockouts, introduce frameshift mutations, and align PduN.

## Notes

https://github.com/agmcfarland/GeneGrouper

### Competing Interest Statement

The authors have declared no competing interest.

