## Supplemental Figures for "Density-based binning of gene clusters to infer function or evolutionary history using GeneGrouper"

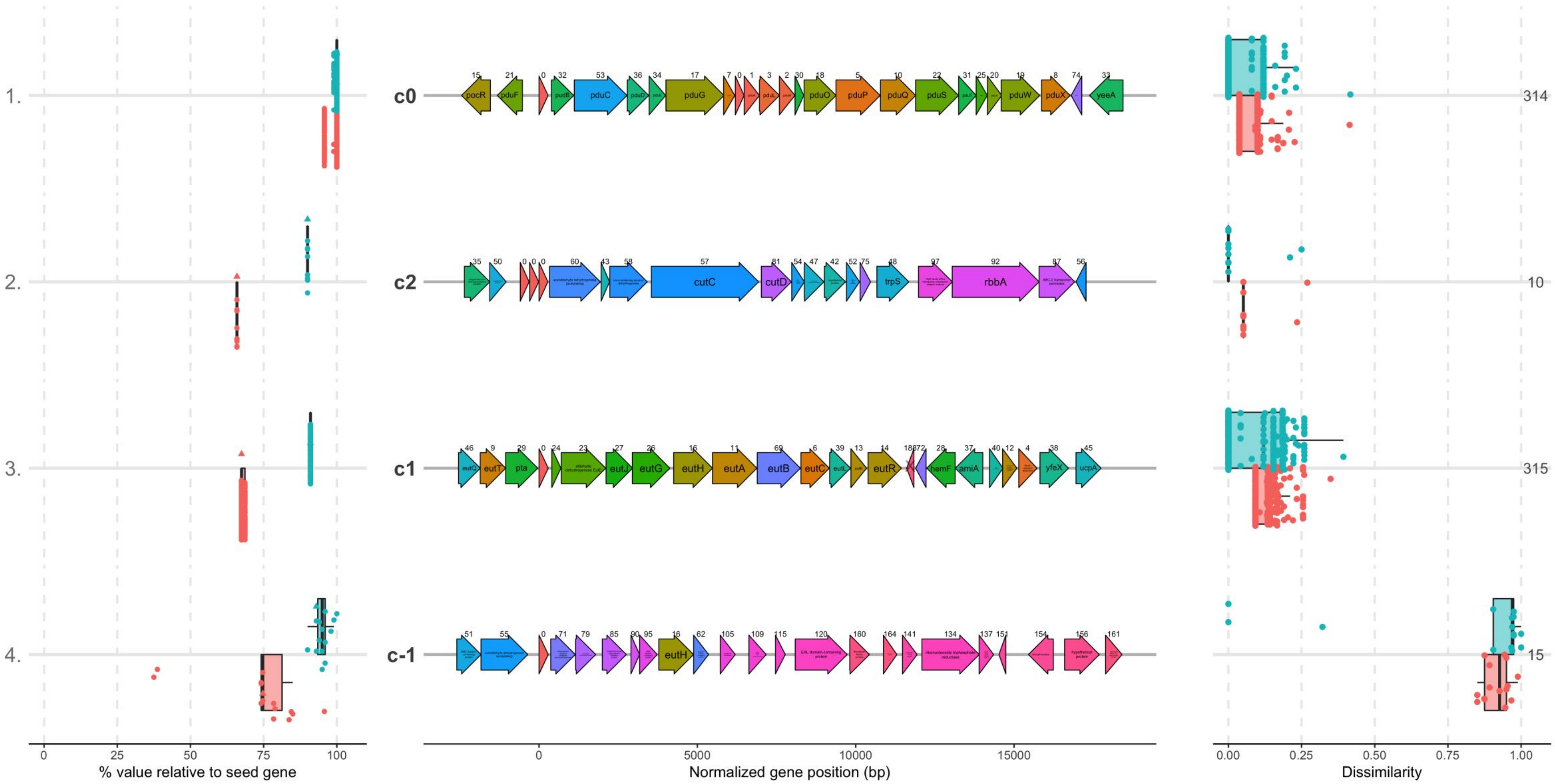

1.
2.
3.
4.
c0
c2
c1
c-1
314
10
315
15
% value relative to seed gene
Normalized gene position (bp)
Dissimilarity
pocR
pduF
pduC
pduG
pduO
pduP
pduQ
pduS
pduW
pduX
yeeA
cutC
cutD
trpS
rbbA
eutQ
eutT
pta
eutJ
eutG
eutH
eutA
eutB
eutC
eutR
hemF
amiA
yfeX
ucpA

A.

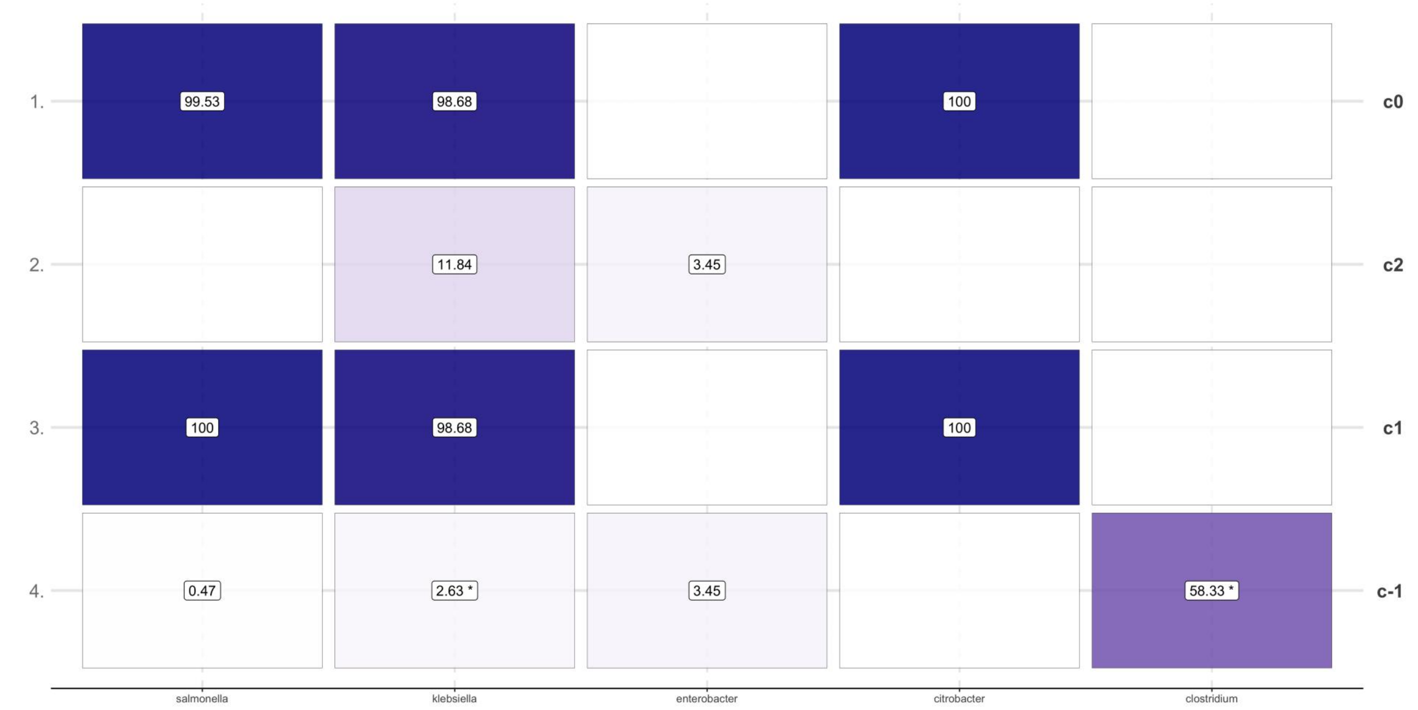
99.53
98.68
100
11.84
3.45
100
98.68
100
0.47
2.63 *
3.45
58.33 *
c0
c2
c1
c-1
salmonella
klebsiella
enterobacter
citrobacter
clostridium

B.

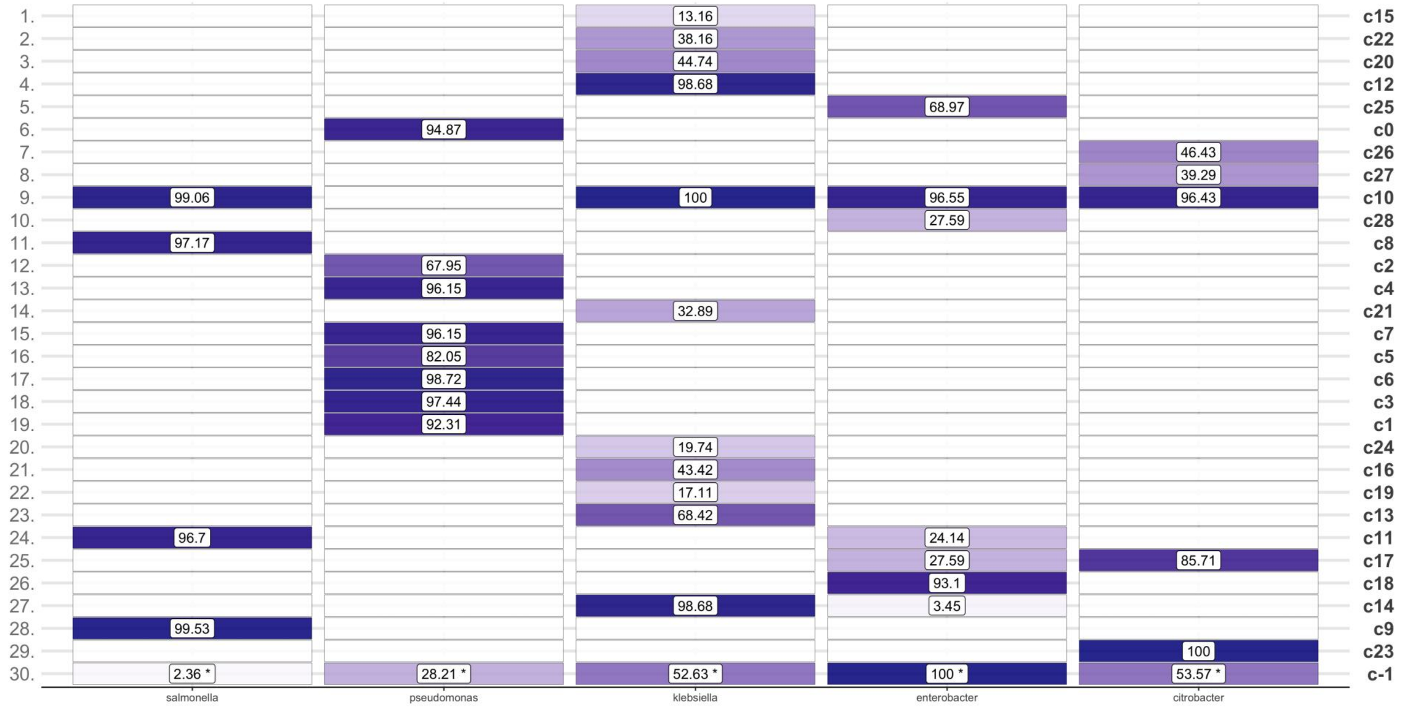
c15
c22
c20
c12
c25
c0
c26
c27
c10
c28
c8
c2
c4
c21
c7
c5
c6
c3
c1
c24
c16
c19
c13
c11
c17
c18
c14
c9
c23
c-1
salmonella
pseudomonas
klebsiella
enterobacter
citrobacter

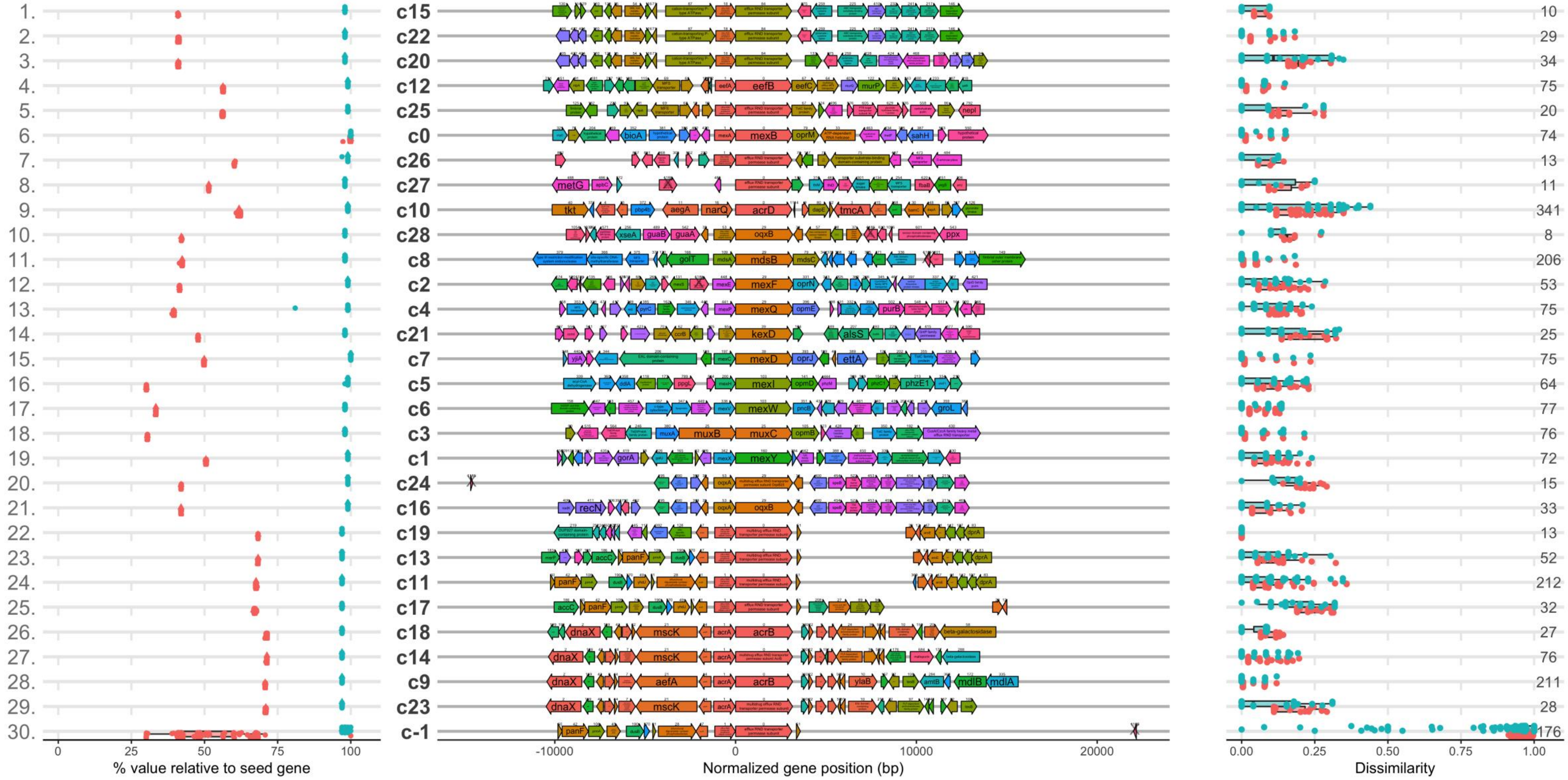

1. c15 10
2. c22 29
3. c20 34
4. c12 75
5. c25 20
6. c0 74
7. c26 13
8. c27 11
9. c10 341
10. c28 8
11. c8 206
12. c2 53
13. c4 75
14. c21 25
15. c7 75
16. c5 64
17. c6 77
18. c3 76
19. c1 72
20. c24 15
21. c16 33
22. c19 13
23. c13 52
24. c11 212
25. c17 32
26. c18 27
27. c14 76
28. c9 211
29. c23 28
30. c-1 176
0 25 50 75 100
% value relative to seed gene
-10000 0 10000 20000
Normalized gene position (bp)
0.00 0.25 0.50 0.75 1.00
Dissimilarity

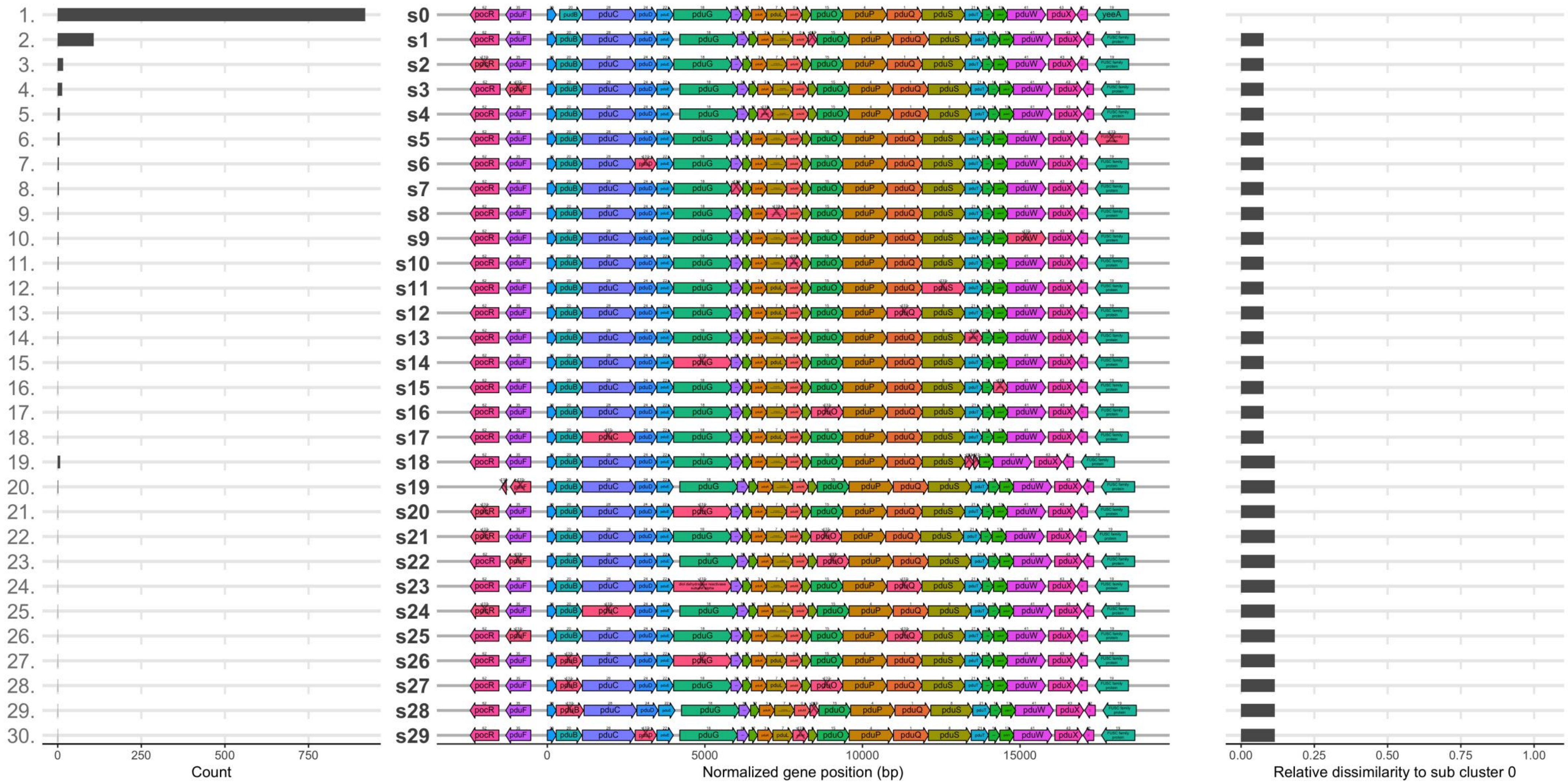
1. 2. 3. 4. 5. 6. 7. 8. 9. 10. 11. 12. 13. 14. 15. 16. 17. 18. 19. 20. 21. 22. 23. 24. 25. 26. 27. 28. 29. 30.
0 250 500 750
Count
s0 s1 s2 s3 s4 s5 s6 s7 s8 s9 s10 s11 s12 s13 s14 s15 s16 s17 s18 s19 s20 s21 s22 s23 s24 s25 s26 s27 s28 s29
pocR pduF pduB pduC pduD pduG pduL pduO pduP pduQ pduS pduW pduX yeeA
0 5000 10000 15000
Normalized gene position (bp)
0.00 0.25 0.50 0.75 1.00
Relative dissimilarity to sub cluster 0

### Alignment of protein:

```
            1        10        20        30        40        50        60        70        80        91
            |--------+---------+---------+---------+---------+---------+---------+---------+---------+|
N_real      MHLARVTGAVVSTQKSPSLIGKKLLLVRRVSADGELPASPTSGDEVAVDSVGAGVGELVLLSGGSSARHVFSGPNEAIDLAVVGIVDTLSC
N_pseudo    MHLARVTGAVVSTQKSPSLIGKSCCWCVGSAPMANSPPRPPPAMKWPWTPSARASANWFCSAAAPAPGTFFPGQMRPSTSPLSAL
Consensus   MHLARVTGAVVSTQKSPSLIGKkcclcrrsaada#lParPppadewawdpsaaasa#lfclaaapaarhfFpGqnraidlalsai......
```

### Alignment of nucleotide:

```
             1        10        20        30        40        50        60        70        80        90        100       110       120       130
             |--------+---------+---------+---------+---------+---------+---------+---------+---------+---------+---------+---------+---------|
PdulN_real   ATGCATCTGGCACGAGTCACGGGCGCGGTTGTCTCCACGCAAAAATCACCTTCTTTGATTGGGAAAAAGCTGCTGCTGGTGCGTCGGGTCAGCGCCGATGGCGAACTCCCCGCCTCGCCCACCTCCGGCG
PdulN_pseudo ATGCATCTGGCACGAGTCACGGGCGCGGTTGTCTCCACGCAAAAATCACCTTCTTTGATTGGGAAAA-GCTGCTGCTGGTGCGTCGGGTCAGCGCCGATGGCGAACTCCCCGCCTCGCCCACCTCCGGCG
Consensus    ATGCATCTGGCACGAGTCACGGGCGCGGTTGTCTCCACGCAAAAATCACCTTCTTTGATTGGGAAAA.GCTGCTGCTGGTGCGTCGGGTCAGCGCCGATGGCGAACTCCCCGCCTCGCCCACCTCCGGCG

             131      140       150       160       170       180       190       200       210       220       230       240       250       260
             |--------+---------+---------+---------+---------+---------+---------+---------+---------+---------+---------+---------+---------|
PdulN_real   ATGAAGTGGCCGTGGACTCCGTCGGCGCGGGCGTCGGCGAACTGGTTTTGCTCAGCGGCGGCTCCAGCGCCAGGCACGTTTTTTCCGGGCCAAATGAGGCCATTGACCTCGCCGTTGTCGGCATTGTAGA
PdulN_pseudo ATGAAGTGGCCGTGGACTCCGTCGGCGCGGGCGTCGGCGAACTGGTTTTGCTCAGCGGCGGCTCCAGCGCCAGGCACGTTTTTTCCGGGCCAAATGAGGCCATCGACCTCGCCGTTGTCGGCATTGTAGA
Consensus    ATGAAGTGGCCGTGGACTCCGTCGGCGCGGGCGTCGGCGAACTGGTTTTGCTCAGCGGCGGCTCCAGCGCCAGGCACGTTTTTTCCGGGCCAAATGAGGCCATcGACCTCGCCGTTGTCGGCATTGTAGA

             261      270  276
             |--------+-----|
PdulN_real   TACGCTTTCGTGTTAA
PdulN_pseudo TACGCTTTCGCGTTAA
Consensus    TACGCTTTCGcGTTAA
```

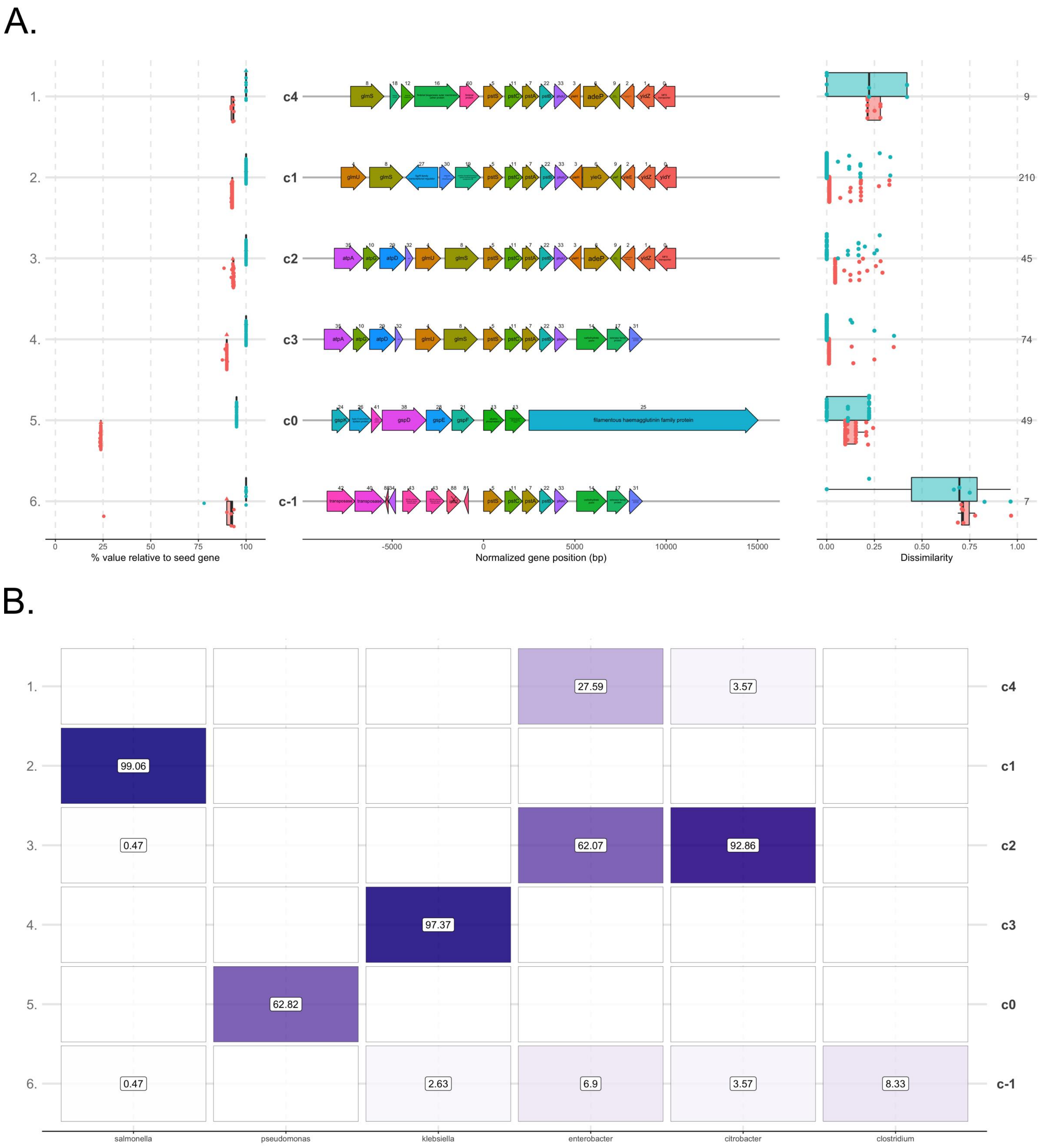
A.
1.
2.
3.
4.
5.
6.
c4
c1
c2
c3
c0
c-1
glmS
pstS
pstC
pstA
adeP
yidZ
glmU
yieG
yidY
atpA
atpD
gspD
gspE
gspF
filamentous haemagglutinin family protein
transposase
9
210
45
74
49
7
0
25
50
75
100
% value relative to seed gene
-5000
0
5000
10000
15000
Normalized gene position (bp)
0.00
0.25
0.50
0.75
1.00
Dissimilarity
B.
27.59
3.57
99.06
0.47
62.07
92.86
97.37
62.82
0.47
2.63
6.9
3.57
8.33
salmonella
pseudomonas
klebsiella
enterobacter
citrobacter
clostridium

A.

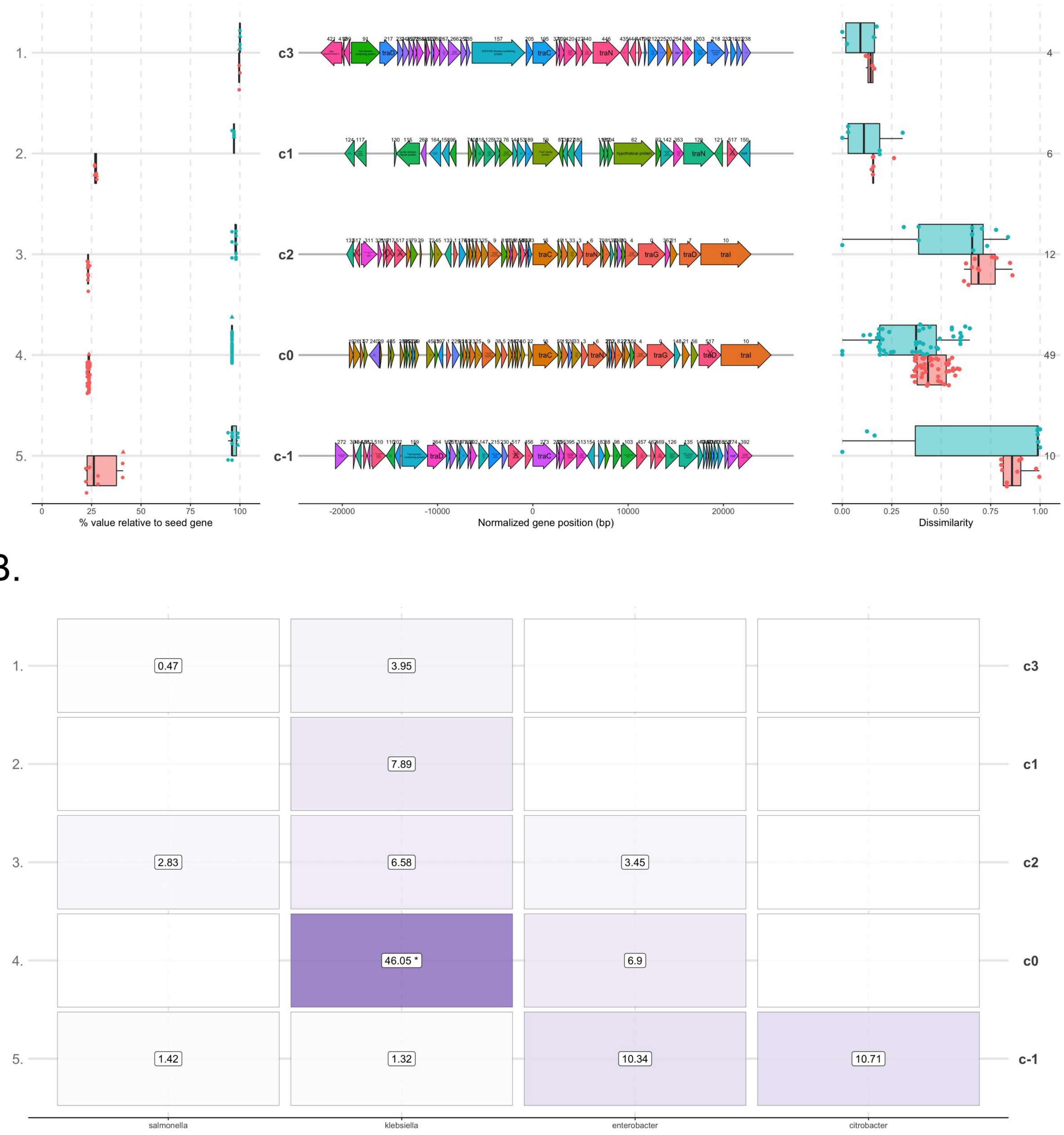

1.
2.
3.
4.
5.
c3
c1
c2
c0
c-1
4
6
12
49
10
% value relative to seed gene
Normalized gene position (bp)
Dissimilarity
traD
traC
traN
traG
traI
hypothetical protein
salmonella
klebsiella
enterobacter
citrobacter
0.47
3.95
7.89
2.83
6.58
3.45
46.05 *
6.9
1.42
1.32
10.34
10.71


B.
