## Supplemental Text for "Density-based binning of gene clusters to infer function or evolutionary history using GeneGrouper"

**Methods**

**Bacterial strain generation**

Modifications to the *pdu* operon of *Salmonella enterica* serovar Typhimurium LT2 were made using the λ Red recombineering method developed by Thomason et al (1). In this method, a *cat/sacB* selectable insert is PCR amplified containing upstream and downstream homologous overhangs corresponding to the target gene locus. The selectable marker is inserted into the desired locus and subsequently knocked out, as in the case of the ΔA ΔJ double knockout strain and ΔN single knockout strain, or replaced with a modified open reading frame, as in the case of the ΔN::N* frameshift mutation. Note that for the ΔN::N* frameshift, a gBlock Gene Fragment from IDT was used. For gene knockouts, 30 bp is left upstream of the downstream open reading frame in order to avoid polar effects. All knockouts and modifications were sequence verified using Sanger sequencing performed by Quintara Biosciences.

**GFP encapsulation assay**

In order to visualize changes to MCP formation and morphology, a GFP encapsulation assay was used as previously described (2). Modified *Salmonella enterica* serovar Typhimurium LT2 were transformed with an inducible fluorescent reporter construct, pBAD33t-ssD-GFPmut2. This plasmid contains an open reading frame that has an N-terminal signal sequence sufficient for targeting the fluorescent reporter, GFPmut2, to the lumen of MCPs. Transformed strains were first streaked from glycerol stocks to single colonies on LB plates supplemented with 34 µg/mL chloramphenicol (incubated at 37 °C for 16 hours after streaking). Single colonies were selected and used to inoculate 5 mL LB liquid cultures supplemented with 34 µg/mL chloramphenicol, which were grown for 16 hours at 37 °C, 225 RPM. These starter cultures were subsequently used to subculture 5 mL LB expression cultures supplemented with 34 µg/mL chloramphenicol, 0.02% (w/v) L-(+)-arabinose, and 0.4% (v/v) 1,2-propanediol. Expression cultures were grown for 6 hours at 37 °C, 225 RPM before imaging.

Cells were imaged using phase contrast and fluorescence microscopy on a Nikon Eclipse Ni-U upright microscope, 100X oil immersion objective, Andor Clara digital camera, and NIS Elements Software (Nikon). Cells were prepared by placing 1.47 µL of the expression culture onto Fisherbrand^TM^ frosted microscope slides and cleaned 22 mm x 22 mm, #1.5 thickness coverslips (VWR Cat# 16004-302). For GFP fluorescence micrographs, the C-FL Endow GFP HYQ bandpass filter was used and images were acquired with an 80 ms exposure. Digital micrographs were processed using ImageJ (3).

**PduN sequence alignments**

The frameshifted *pduN* (N*) pseudogene was downloaded from the National Center for Biotechnology Information (NCBI) website (reference sequence NC_004631.1) and compared to the *pduN* sequence from *Salmonella enterica* serovar Typhimurium. The translated sequences were produced using the Expert Protein Analysis System (ExPASy) Translate web tool (4), and alignments were created and exported using the MultAlin webtool (5).
